# The Impact of Pathway Database Choice on Statistical Enrichment Analysis and Predictive Modeling

**DOI:** 10.1101/654442

**Authors:** Sarah Mubeen, Charles Tapley Hoyt, Andre Gemünd, Martin Hofmann-Apitius, Holger Fröhlich, Daniel Domingo-Fernández

## Abstract

**Background:** Pathway-centric approaches are widely used to interpret and contextualize -*omics* data. However, databases contain different representations of the same biological pathway, which may lead to different results of statistical enrichment analysis and predictive models in the context of precision medicine.

**Results:** We have performed an in-depth benchmarking of the impact of pathway database choice on statistical enrichment analysis and predictive modeling. We analyzed five cancer datasets using three major pathway databases and developed an approach to merge several databases into a single integrative database: MPath. Our results show that equivalent pathways from different databases yield disparate results in statistical enrichment analysis. Moreover, we observed a significant dataset-dependent impact on performance of machine learning models on different prediction tasks. Further, MPath significantly improved prediction performance and reduced the variance of prediction performances in some cases. At the same time, MPath yielded more consistent and biologically plausible results in the statistical enrichment analyses. Finally, we implemented a software package designed to make our comparative analysis with these and additional databases fully reproducible and to facilitate the update of our integrative pathway resource in the future.

**Conclusion:** This benchmarking study demonstrates that pathway database choice can influence the results of statistical enrichment analysis and prediction modeling. Therefore, we recommend the use of multiple pathway databases or the use of integrative databases.

## 1. Introduction

As fundamental interactions within complex biological systems have been discovered in experimental biology labs, they have often been assembled into computable pathway representations. Because they have proven immensely useful in the analysis and interpretation of -*omics* data when coupled with algorithmic approaches (e.g., gene set enrichment analysis (GSEA)), academic and commercial groups have generated and maintained a comprehensive set of databases during the last 15 years (Bader *et al*., 2006). Examples include KEGG, Reactome, WikiPathways, NCIPathways and Pathway Commons (Kanehisa *et al*., 2016; Fabregat *et al.*, 2018; Slenter *et al.*, 2017; Schaefer *et al.*, 2008; Cerami *et al.*, 2011).

However, they tend to differ in the average number of pathways they contain, the average number of proteins per pathway, the types of biochemical interactions they incorporate, ands the subcategories of pathways that they provide (e.g., signal transduction, genetic interaction and metabolic) (Kirouac *et al.*, 2012; Türei et al., 2016). Pathways are often also described at varying levels of detail, with diverse data types and with loosely defined boundaries (Domingo-Fernández *et al.*, 2018). Nonetheless, most pathway analyses are still conducted exclusively by employing a single database, often chosen in part by researchers’ preferences or previous experiences (e.g., bias towards a database previously yielding good results, ease of use of a particular database) (Table 1). Notably, the selection of a suitable pathway database depends on the actual biological context that is investigated, yet KEGG remains severely over-represented in published -*omics* studies. This raises concerns and motivates the consideration of multiple pathway databases or, preferably, an integration over several pathways resources.

**Table 1.** Number of publications citing major pathway resources for pathway enrichment in PubMed Central (PMC), 2019. To develop an estimate on the number of publications using several pathway databases for pathway enrichment, SCAIView (http://academia.scaiview.com/academia; indexed on 01/03/2019) was used to conduct the following query using the PMC corpus: “<pathway resource>” AND “pathway enrichment”.

| Type | Pathway Resource | Publications |
| --- | --- | --- |
| Primary | KEGG | 27,713 |
|  | Reactome | 3,765 |
|  | WikiPathways | 651 |
| Integrative | MSigDB | 2,892 |
|  | ConsensusPathDB | 339 |
|  | Pathway Commons | 1640 |

Several integrative resources have been developed, including meta-databases (e.g., Pathway Commons; Cerami *et al.*, 2011, MSigDB; Liberzon *et al.*, 2015, and ConsensusPathDB; Kamburov *et al.*, 2008) that enable pathway exploration in their corresponding web applications and integrative software tools (e.g., graphite; Sales *et al.*, 2018, PathMe; Domingo-Fernández *et al*, 2019, and OmniPath; Türei *et al.*, 2016) designed to enable bioinformatics analyses. By consolidating pathway databases, these resources have attempted to summarize major reference points in the existing knowledge and demonstrate how data contained in one resource can be complemented by data contained in others. Thus, through their usage, the biomedical community has benefitted from comprehensive overviews of pathway landscapes which can then make for more robust resources highly suited for analytic usage.

The typical approach to combine pathway information with -*omics* data is via statistical enrichment analysis, also known as pathway enrichment. The task of navigating through the continuously developing variants of enrichment methods has been undertaken by several recent studies which benchmarked the performance of these techniques (Bayerlová *et al.*, 2015; Ihnatova *et al*., 2018; Lim *et al*., 2018) and guide users on the choice for their analyses (Fabris *et al.*, 2019; Reimand *et al.*, 2019). While Bateman *et al*. (2014) examined the impact of choice of different subsets of MSigDB on GSEA analysis, it remains unclear what broader impact an integrative pathway meta-database would have for statistical enrichment analysis. Additionally, the overlap of pathways within the same integrative database can induce biases (Liberzon *et al.*, 2015), specifically when conducting multiple testing correction via the popular Benjamini-Hochberg method (Benjamini and Hochberg, 1995) that supposes independence of statistical tests. This issue is of particular concern for large-scale meta-databases such as MSigDB.

The aim of this work is to systematically investigate the influence of alternative representations of the same biological pathway (e.g. in KEGG, Reactome and WikiPathways) on the results of statistical enrichment analysis via three common methods: the hypergeometric test, GSEA and Signaling Pathway Impact Analysis (SPIA) (Fisher, 1992; Subramanian *et al*., 2005; Tarca *et al*., 2008) using five The Cancer Genome Atlas (TCGA) datasets (Weinstein *et al.*, 2013). In addition, we also show that pathway activity based patient classification and survival analysis via single sample GSEA (ssGSEA; Barbie *et al.*, 2009) can be impacted by the choice of pathway resource in some cases. As a solution, we propose to integrate different pathway resources via a method where semantically analogous pathways across databases (e.g., “Notch signaling pathway” in KEGG and “Signaling by NOTCH” pathway in Reactome) are combined. This approach exploits the pathway mappings and harmonized pathway representations described in our previous work (Domingo-Fernández *et al*., 2018; Domingo-Fernández *et al*., 2019). We demonstrate that when aided by our integrative pathway database, it is possible to better capture expected disease biology than with individual resources, and to sometimes obtain better predictions of clinical endpoints. Our entire analytic pipeline is implemented in a reusable Python package (`pathway_forte`; see Methods) to facilitate reproducing the results with other databases or datasets in the future.

## 2. Results

The results of the benchmarking study have been divided into two subsections for each of the pathway methods described above. We first compared the effects of database selection on the results of functional pathway enrichment methods. In the following subsection, we benchmarked the performance of the pathway resources on the various machine learning classification tasks conducted.

### 2.1. Benchmarking the impact on enrichment methods

#### 2.1.1. Over-representation analysis

As illustrated by our results, pathway analogs from different pathway databases in several cases showed clearly significant rank differences (Figure 1). These differences were most pronounced between Reactome and WikiPathways. For example, while the “Thyroxine Biosynthesis” pathway was highly statistically significant (*q*-value < 0.01) in the LIHC dataset for Reactome, its analogs in WikiPathways (i.e., “Thyroxine (Thyroid Hormone) Production”) and KEGG (i.e., “Thyroid Hormone Synthesis”) were not. However, the pathway was found to be significantly enriched in MPath. Such differences were similarly observed for the “Notch signalling” pathway in the PRAD dataset, in which the pathway was highly statistically significant (*q*-value < 0.01) for Reactome and MPath, but showed no statistical significance for KEGG and WikiPathways. Similar cases were systematically observed for additional pathway analogs and super pathways, demonstrating that marked differences in rankings can arise depending on the database used.

**Figure 1:**
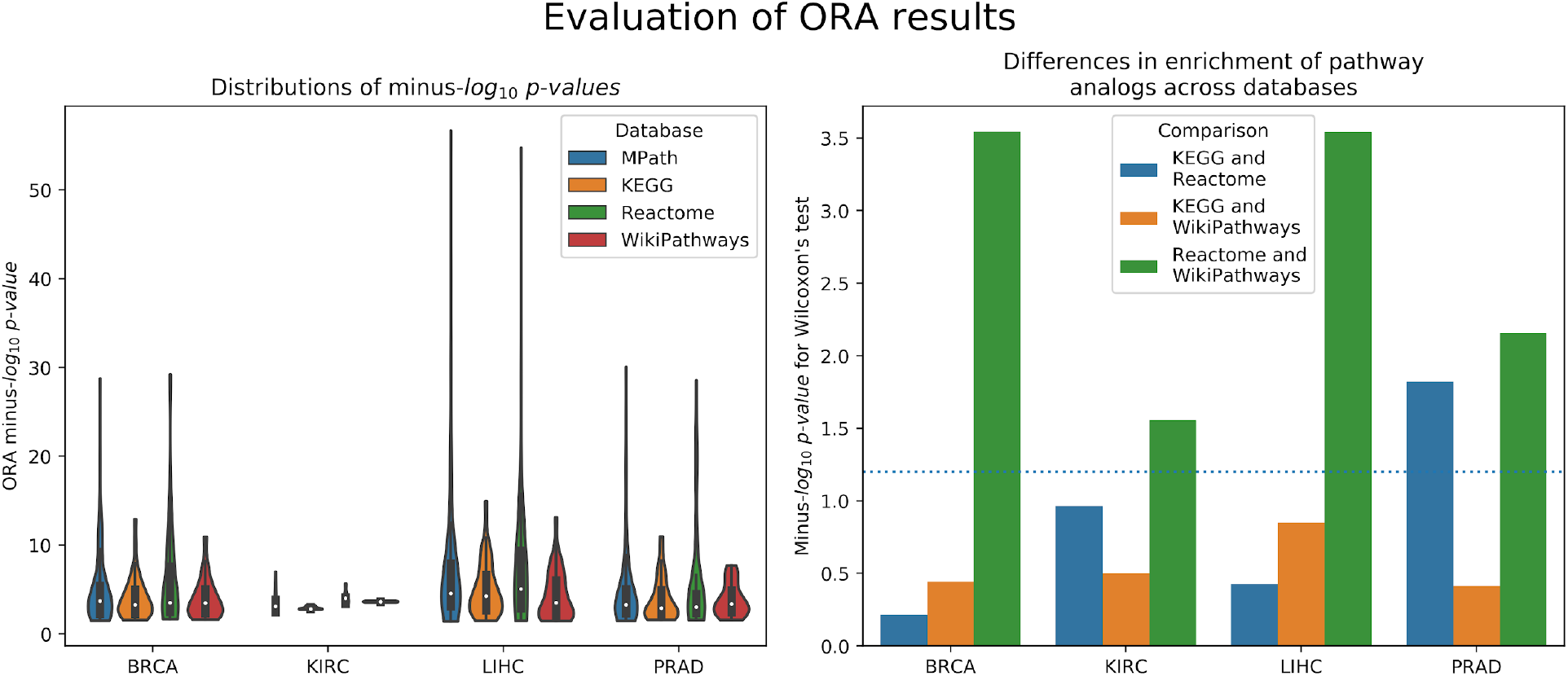
(left) Distribution of raw *p-*values of pathway analogs across databases (ORA). (right) Significance of average rank differences of pathway analogs across pairwise database comparisons.

#### 2.1.2. Gene Set Enrichment Analysis

Similar to ORA, GSEA showed significant differences between pathway analogs across databases in several cases (Figure 2). These differences were most pronounced between KEGG and WikiPathways in the KIRC and LIHC datasets, and between KEGG and Reactome in the BRCA and PRAD datasets. Since GSEA calculates the observed direction of regulation (e.g., over/under-expressed) of each pathway, we also examined whether super pathways or pathway analogs exhibited opposite signs in their Normalized Enrichment Scores (NES) (e.g., one pathway is over-expressed while its equivalent pair is under-expressed). As an illustration, GSEA results of the LIHC dataset revealed the contradiction that the “DNA replication” pathway, one of 26 super pathways, was over-expressed according to Reactome, and under-expressed according to KEGG and WikiPathways, though the pathway was not statistically significant for any of these databases. However, the merged “DNA replication” pathway in MPath appeared as significantly under-expressed. Similarly, in the BRCA dataset, the WikiPathways definition of the “Notch signaling” and “Hedgehog signaling” pathways were significantly over-expressed, while the KEGG and Reactome definitions were insignificantly over-expressed. Interestingly, both the merged “Notch signaling” and merged “Hedgehog signaling” pathways appeared as significantly under-expressed (*q* < 0.05) in MPath.

**Figure 2:**
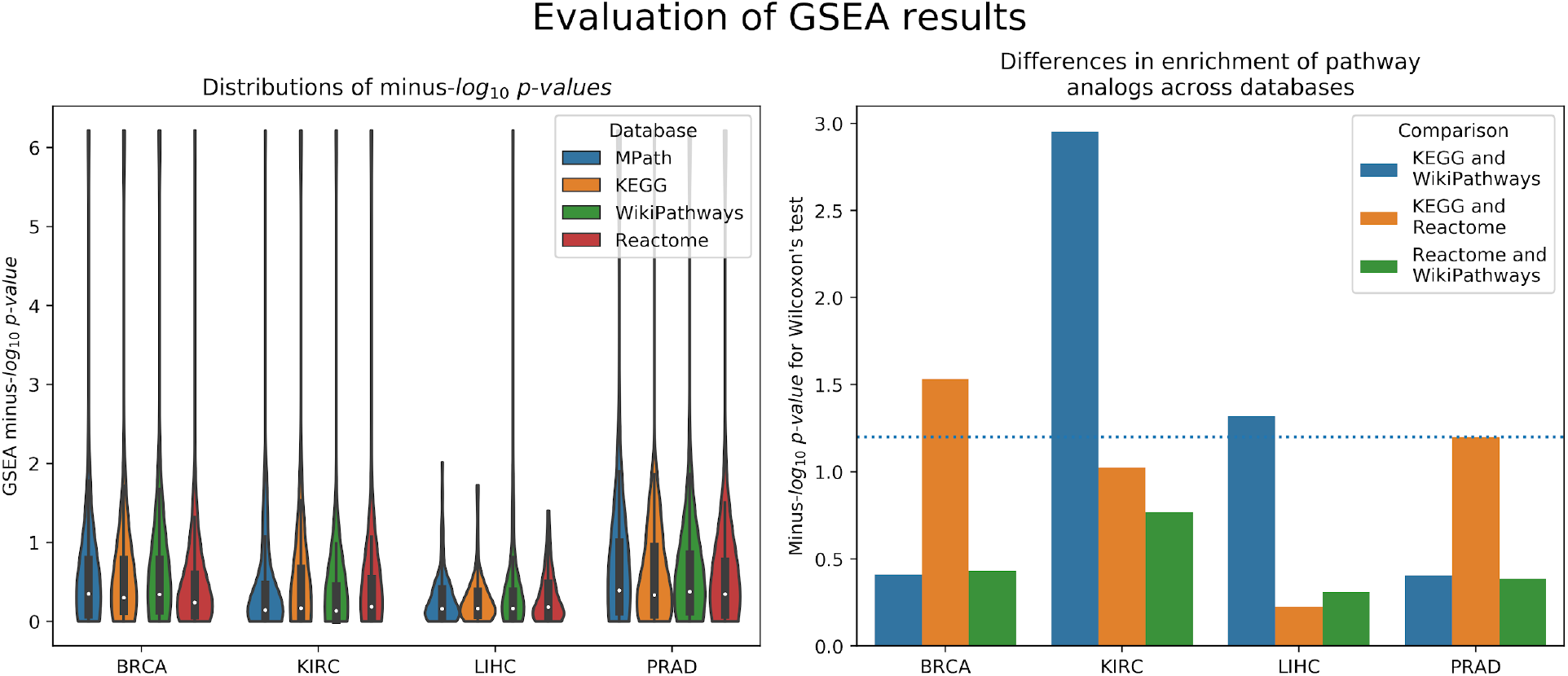
(left) Distribution of raw *p-*values of pathway analogs across databases (GSEA). (right) Significance of average rank differences of pathway analogs across pairwise database comparisons.

#### 2.1.3. Signaling Pathway Impact Analysis

The final of the three statistical enrichment analyses conducted revealed further differences between pathway analogs across databases. As expected, differences in the results of analogous pathways were exacerbated on topology-based methods compared with ORA and GSEA, as these latter methods do not consider pathway topology (i.e., incorporation of pathway topology introduces one extra level of complexity leading to higher variability) (Figure 3). Beyond a cursory inspection of the statistical results, we also investigated the concordance of the direction of change of pathway activity (i.e., activation or inhibition) for equivalent pathways. We found that for two database (i.e., LIHC and KIRC), the direction of change was inconsistently reported for the “TGF beta signaling” pathway, depending on the database used (i.e., the KEGG representation was activated and the WikiPathways one inhibited). A similar effect was observed in the “Estrogen signaling pathway”, found to be inhibited in KEGG and activated in WikiPathways in the LIHC dataset. The merging of equivalent pathway networks resulted in the observation of inhibition for both the “TGF beta signaling” and “Estrogen signaling” pathways in MPath results.

**Figure 3:**
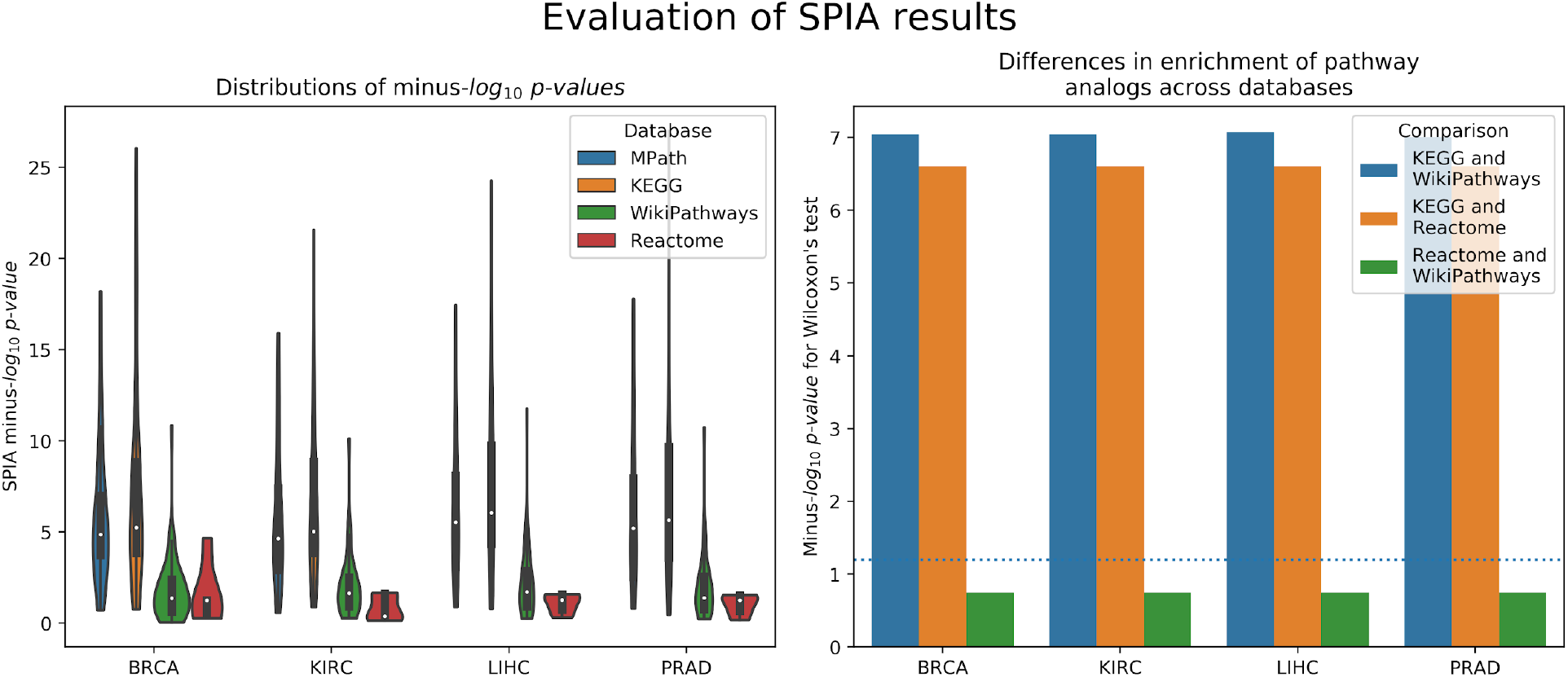
(left) Distribution of raw *p-*values of pathway analogs across databases (SPIA). (right) Significance of average rank differences of pathway analogs across pairwise database comparisons.

### 2.2. Benchmarking the impact on predictive modeling

#### 2.2.1. Prediction of tumor vs. normal

We compared the prediction performance of an elastic net penalized logistic regression classifier to discriminate normal from cancer samples based on their pathway activity profiles. The cross-validated prediction performance was measured via the area under ROC curve (AUC) (see the corresponding methods section). Our results indicated no overall significant effect of the choice of pathway database on model prediction performance (*p* = 0.5, ANOVA F-test, Figure 4). However, Tukey’s post-hoc analysis revealed a significant improvement of MPath based pathway signatures compared to WikiPathways for BRCA (95% CI: 0.97% −2.3%).

**Figure 4.**
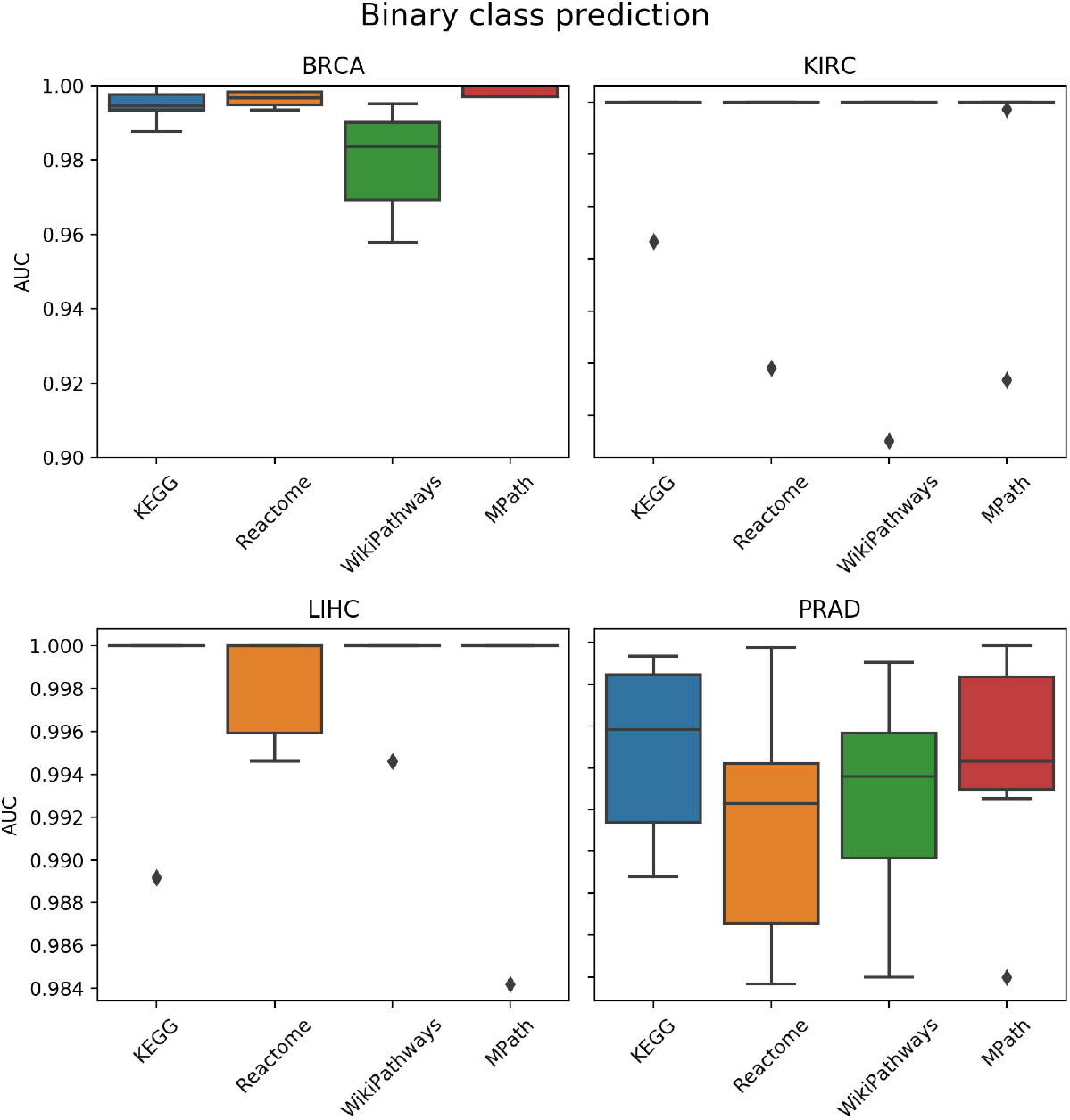
Comparison of prediction performance of an elastic net classifier (tumor vs. normal) using ssGSEA based pathway activity profiles computed from different resources. Each boxplot shows the distribution of the AUCs over 10 repeats of the 10-fold cross-validation procedure.

#### 2.2.2. Prediction of tumor subtype

We next compared the prediction performances of a multi-class classifier predicting known tumor subtypes of BRCA and PRAD using ssGSEA based pathway activity profiles. Figure 5 demonstrated no overall significant effect of the choice of pathway database (*p* = 0.16, ANOVA F-test). Nonetheless, Tukey’s post-hoc analysis again demonstrated a significant benefit of MPath based pathway signatures compared to KEGG (95% CI: 0.76% - 7.8%) for BRCA.

**Figure 5.**
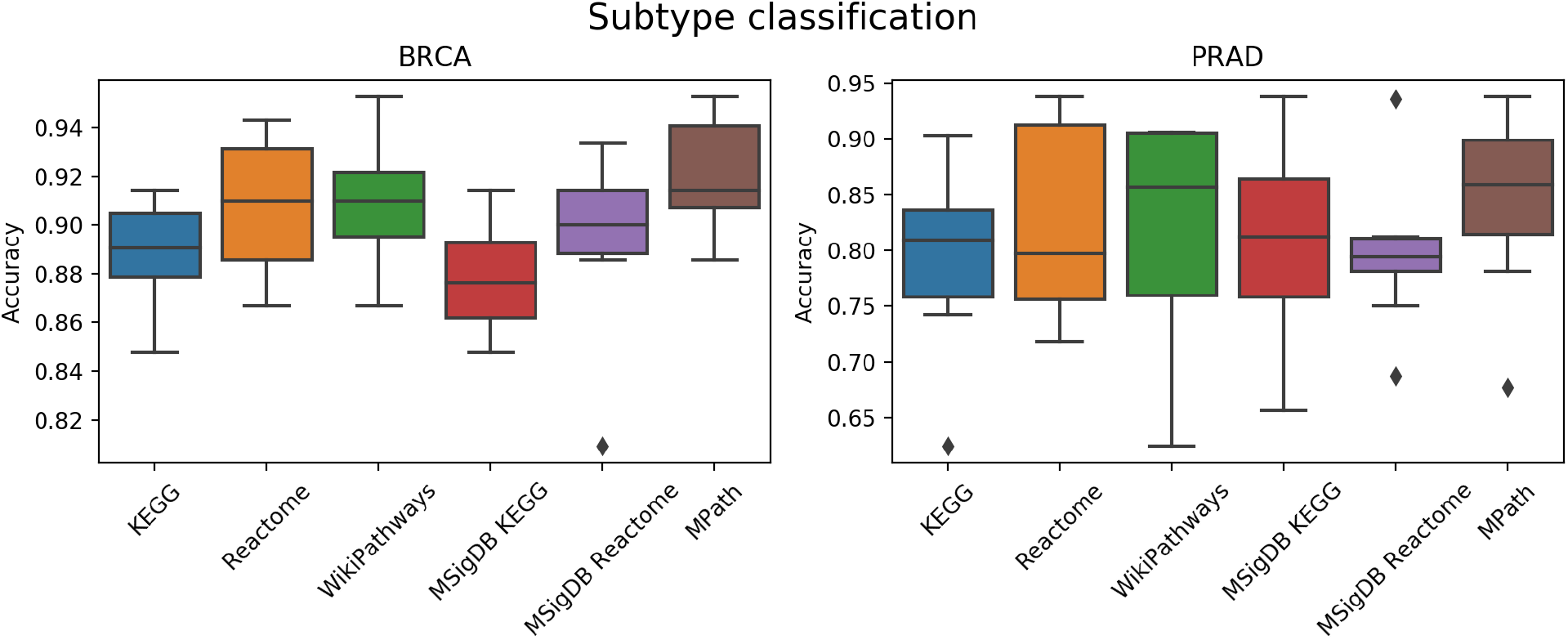
Comparison of prediction performance of an elastic net classifier (BRCA and PRAD subtypes) using ssGSEA based pathway activity profiles computed from different resources. Each boxplot shows the distribution of the AUCs over 10 repeats of the 10-fold cross-validation procedure.

#### 2.2.3. Prediction of overall survival

As a next step, we compared the prediction performance of an elastic net penalized Cox regression model for overall survival using ssGSEA based pathway activity profiles derived from different resources. As indicated in Figure 6, no overall significant effect of the actually used pathway database could be observed (*p* = 0.28, ANOVA F-test), but for specific datasets Tukey’s post-hoc analysis again revealed significant differences, more specifically of MSigDB KEGG vs Reactome for BRCA (95% CI: 0.11% - 9.97%). A limiting factor of this analysis is the fact that overall survival can generally only be predicted slightly above chance level (c-indices range between 55 - 60%) based on gene expression alone, which is in agreement to the literature (Fröhlich, 2014; Mayr and Schmid, 2014; Van Wieringen *et al.*, 2009; Zhang *et al.*, 2018).

**Figure 6.**
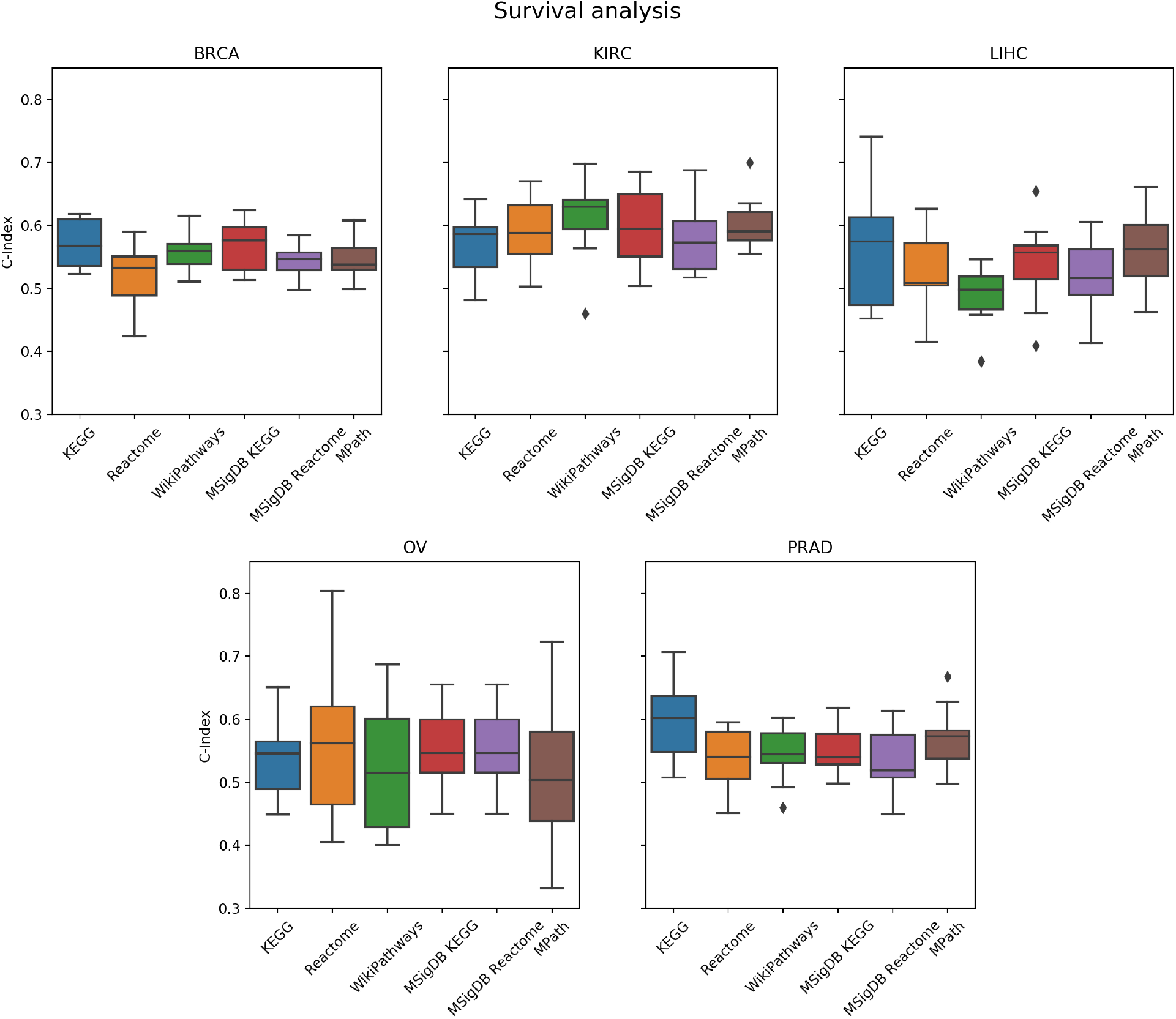
Comparison of prediction performance of an elastic net penalized Cox regression model (overall survival) using ssGSEA based pathway activity profiles computed from different resources. Each boxplot shows the distribution of the AUCs over 10 repeats of the 10-fold cross-validation procedure.

## 3. Discussion

In this work, we presented a comprehensive comparative study of pathway databases based on functional enrichment and predictive modeling. We have shown that the choice of pathway database can significantly influence the results of statistical enrichment, which raises concerns about the typical lack of consideration that is given to the choice of pathway resource in many gene expression studies. This finding was specifically pronounced for SPIA, because SPIA is a topology based enrichment approach and therefore expected to be most sensitive to the actual definition of a pathway. At the same time, we observed that an integrative pathway resource (MPath) led to more biologically consistent results and in some cases, improved prediction performance.

However, generating a merged dataset such as MPath is non-trivial. We purposely restricted this study to three major pathway databases because of the availability of inter-database pathway mappings, harmonized gene sets and pathway networks from our previous work which enabled conducting objective database comparisons. However, this effort implies further curation and harmonization with the incorporation of each additional pathway database into MPath for future benchmarking studies.

Our strategy to build MPath is one of many possible approaches to integrate pathway knowledge from multiple databases. Although alternative meta-databases such as Pathway Commons and MSigDB do exist, the novelty of this work lies in the usage of mappings and harmonized pathway representations for generating a merged dataset. While we have presented MPath as one possible integrative approach, alternative meta-databases may be used, but would require that researchers ensure that the meta-databases’ contents are continuously updated (Wadi *et al.*, 2016).

Our developed mapping strategy between different graph representations of analogous pathways enabled us to objectively compare pathway enrichment results that otherwise would have been conducted manually and subjectively. Furthermore, they allowed us to generate super pathways inspired by previous approaches that have shown the benefit of merging similar pathway representations (Stoney *et al*., 2018; Vivar *et al*., 2013; Doderer *et al*., 2012; Belinky *et al*., 2015). In this case, this was made possible by the fully harmonized gene sets and networks generated by our previous work, ComPath and PathMe.

One of the limitations of this work is that we restricted the analysis to five cancer datasets from TCGA and we did not expand it to other conditions besides cancer. The use of this disease area was mainly driven by the availability of data and the corresponding possibilities to draw statistically valid conclusions. However, we acknowledge the fact that data from other disease areas may result in different findings. More specifically, we believe that a similar benchmarking study based on data from disease conditions with an unknown pathophysiology (e.g., neurological disorders) may have yield even more pronounced differences between pathway resources. Additionally, further techniques for gene expression based pathway activity scoring could be incorporated such as Pathifier or SAS (Drier *et al.*, 2013; Lim *et al.*, 2016).

## 4. Conclusion

In this study, we have systematically investigated the influence of the choice of pathway database on various techniques for functional pathway enrichment and different predictive modeling tasks. Collectively, this study has made three contributions: i) we have shown that there are differences based on the choice of pathway database, ii) we have shown that these differences can be mitigated by using integrative databases and iii) we have implemented a software to facilitate similar benchmarking studies in the future and to re-apply our pathway integration strategy to other resources.

## 5. Methods

In the first two subsections, we describe the pathway resources and the clinical and genomic datasets we used in benchmarking. The following sections then outline the statistical enrichment analysis and predictive modeling conducted in this study. Finally, in the last two subsections, we describe the statistical methods and the software implemented to conduct the benchmarking.

### 5.1. Pathway databases

#### 5.1.1. Selection criteria

Numerous viable pathway databases have been made available to infer biologically relevant pathway activity (Bader *et al.*, 2006). In this work, we systematically compared three major ones (i.e., KEGG, Reactome and WikiPathways) as the subset of databases to benchmark. The rationale for the inclusion of these databases was two-fold: firstly, these databases are open sourced, well-established and highly-cited in studies investigating pathways associated with variable gene expression patterns in different sets of conditions (Table 1). Secondly, we expected distinctions between these databases to be strong enough to observe variable results of enrichment analysis and patient classification, yet these databases also contain a reasonable number of equivalent pathways such that objective comparisons could be made, as outlined in our previous work (Domingo-Fernández *et al*., 2018).

#### 5.1.2. Data retrieval and processing

In order to systematically compare results yielded by different databases, we retrieved the contents of KEGG, Reactome and WikiPathways using ComPath (Domingo-Fernández *et al*., 2018) and converted it into the Gene Matrix Transposed (GMT) file format. Generated networks encoded in Biological Expression Language (BEL; Slater, 2014) were retrieved using PathMe (Domingo-Fernández *et al*., 2019).

To test the potential utility of an integrative pathway resource, we used equivalent pathways across the three databases that were manually curated in our previous work (Domingo-Fernández *et al*., 2018) (see our earlier publication for further details). In the following, we call these “pathways analogs” or “equivalent pathways” (Figure 7a), while we call a pathway found as analogous across all KEGG, Reactome as well as WikiPathways a “super pathway”.

**Figure 7.**
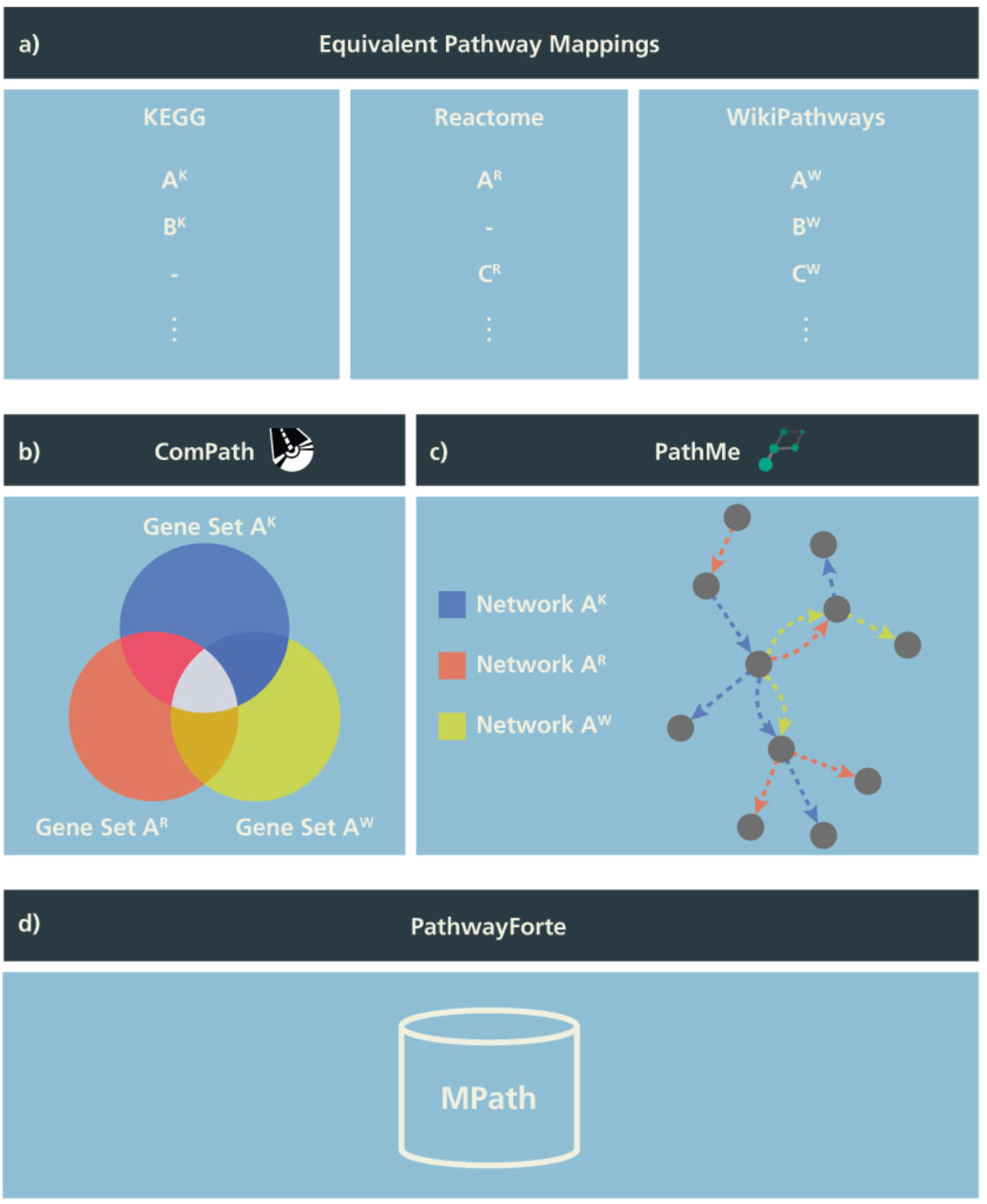
Schema illustrating the generation of MPath. The curated pathway mapping catalog is depicted in **a)** which links equivalent pathways from different resources. Pathways that are shared across two resources are referred to as pathway analogs (i.e., Pathway A in Reactome and Pathway A’ in KEGG) and pathways that are shared across all three resources are referred to as “super pathways” (i.e., Pathway A in KEGG, Pathway A’ in Reactome and Pathway A’’ in WikiPathways). **b)** Using these mappings, gene sets of equivalent pathways from different resources can be combined, ensuring key molecular players from the different resources are included. **c)** Similarly, network representations of the pathways can be overlaid to generate more comprehensive pathways. **d)** Finally, both the combined gene sets and networks representations are included in MPath. Note that pathways that are exclusive to a single database are included in MPath unchanged.

In a second step, we merged equivalent pathways by taking the graph union with respect to contained genes and interactions (Figure 7b and c). We have also described this step in more detail in our earlier work (Domingo-Fernández *et al*., 2019).

The set union of KEGG, Reactome and WikiPathways, while taking into account pathway equivalence, gave rise to an integrative resource to which we refer as *MPath* (Figure 7d). By merging equivalent pathways, MPath contains a fewer number of pathways than the sum of all pathways from all primary resources. In total, MPath contains 2896 pathways of which 238 are derived from KEGG, 2119 from Reactome and 409 from WikiPathways, while another 129 pathways are pathway analogs and 26 are super pathways.

We next compared the latest versions of pathway gene sets from KEGG, Reactome, WikiPathways and MPath with pathway gene sets from MSigDB, a highly cited integrative pathway database containing older versions of the KEGG and Reactome gene sets (ziberzon *et al.*, 2015). We downloaded KEGG and Reactome gene sets from the curated gene set (C2) collection of MSigDB (URL: http://software.broadinstitute.org/gsea/msigdb/collections.jsp#C2; version 6.2; July 2018). Detailed statistics on the number of pathways from each resource are presented in Table 2.

**Table 2.** Statistics of the six pathway resources used in this work. The statistics for KEGG, Reactome, WikiPathways and MPath correspond to the number of pathways retrieved from ComPath on the 28^th^ of February, 2019. Gene sets from MSigDB correspond to the 6.2 release (July 2018).

| Pathway Resource | Pathways |
| --- | --- |
| KEGG | 330 |
| Reactome | 2,219 |
| WikiPathways | 511 |
| MPath | 2,896 |
| MSigDB KEGG | 186 |
| MSigDB Reactome | 674 |

### 5.2. Clinical and genomic data

We used five widely-used datasets acquired from TCGA (Weinstein *et al.*, 2013), a cancer genomics project that has catalogued molecular and clinical information for normal and tumor samples (Table 3). TCGA data were retrieved through the Genomic Data Commons (GDC; https://gdc.cancer.gov) portal and cBioportal (https://www.cbioportal.org) on the 14th of March, 2019. RNA-seq gene expression data subjected to an mRNA quantification analysis pipeline for BRCA, KIRC, LIHC, OV and PRAD TCGA datasets were queried, downloaded and prepared from the GDC through the R/Bioconductor package, TCGAbiolinks (R version: 3.5.2; TCGAbiolinks version: 2.10.3), (Colaprico, *et al.*, 2015). The data were preprocessed as follows: gene expression was quantified by the number of reads aligned to each gene and read counts were measured using HTSeq and normalized using Fragments Per Kilobase of transcript per Million mapped reads upper quartile (FPKM-UQ). HTSeq raw read counts also subject to the GDC pipeline were similarly queried, downloaded and prepared with TCGAbiolinks. Read count data downloaded for the BRCA, KIRC, LIHC and PRAD datasets were processed to remove identical entries while unique measurements of identical genes were averaged. The differential gene expression analysis of cancer versus normal samples was performed using the R/Bioconductor package, DESeq2 (version 1.22.2). Genes with adjusted *p*-value < 5% were considered significantly dysregulated. For all downloaded data, gene identifiers were mapped to HGNC gene symbols (Povey *et al.*, 2001), where possible. To obtain additional information on the survival status and time to death, or censored survival times of patients, patient identifiers in the TCGA datasets were mapped to their equivalent identifiers in cBioPortal. Additionally, cancer subtype classifications or the PRAD and BRCA datasets were retrieved from the GDC. We would like to note that although there are other cohorts available (e.g., COAD and STAD) containing all of these modalities, we did not include them in this analysis because of the limited number of samples they contain (i.e., less than 300 patients). Detailed statistics of all five datasets are presented in Table 3.

**Table 3.** Statistics of the five TCGA cancer datasets used in this work. The statistics correspond to those retrieved from the GDC portal and cBioportal on the 14th of March, 2019. Longitudinal statistics of survival data are presented in **Supplementary Figure 1.**

| Cancer type | TCGA | Tumor | Normal | Surviving | Deceased |
| --- | --- | --- | --- | --- | --- |

|  | abbreviation | samples | samples | patients | patients |
| --- | --- | --- | --- | --- | --- |
| Breast invasive carcinoma | BRCA | 1102 | 113 | 946 | 153 |
| Kidney renal clear cell carcinoma | KIRC | 538 | 72 | 365 | 173 |
| Liver hepatocellular carcinoma | LIHC | 371 | 50 | 240 | 130 |
| Prostate adenocarcinoma | PRAD | 498 | 52 | 498 | 10 |
| Ovarian cancer | OV | 374 | 0 | 143 | 229 |

### 5.3. Pathway enrichment methods

In this subsection, we describe three different classes of pathway enrichment methods that we tested: i) statistical over-representation analysis (ORA), ii) functional class scoring (FCS) and iii) pathway topology (PT) based enrichment (Figure 8) (Khatri *et al*., 2012; García-Campos *et al*., 2015; Fabris *et al*., 2019).

**Figure 8.**
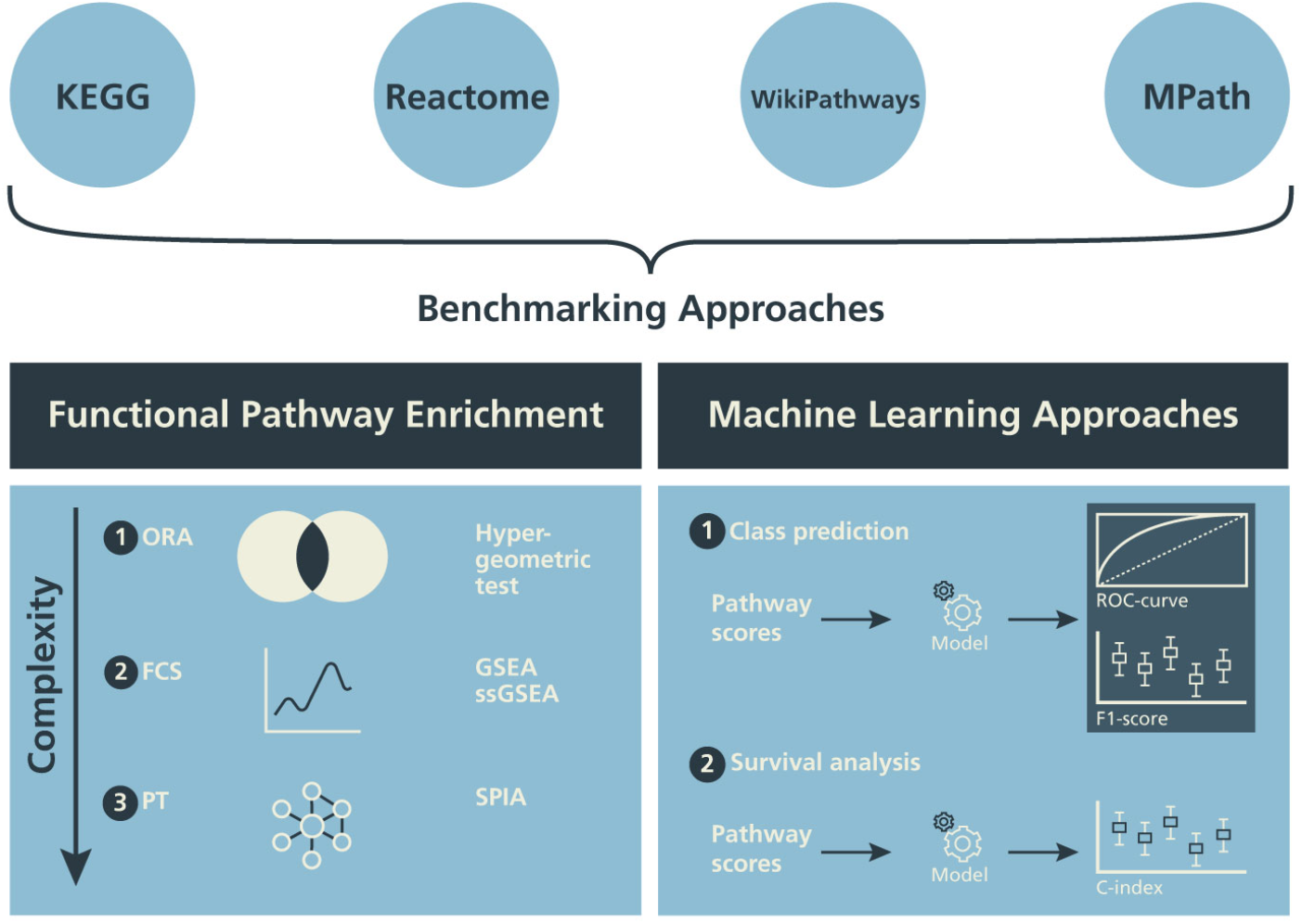
Design of the benchmarking schema. The influence of alternative pathway databases on the results of statistical pathway enrichment (a) and machine learning classification tasks (b) are compared.

#### 5.3.1. Over-representation analysis

We conducted pathway enrichment using genes that exhibited a *q*-value < 0.05 using a one-sided Fisher’s exact test (Fisher, 1992) for each of the pathways in all pathway databases. We consider a pathway to be significantly enriched if its *q*-value is smaller than 0.05 after applying multiple hypothesis testing correction with the Benjamini–Yekutieli method under dependency (Benjamini and Yekutieli, 2001).

#### 5.3.2. Functional class scoring methods

We selected GSEA, one of the most commonly used FCS methods (Subramanian *et al*., 2005). We performed GSEA with the Python package, GSEApy (version 0.9.12; https://github.com/zqfang/gseapy), using normalized RNA-seq expression quantifications (FPKM-UQ) obtained for the BRCA, KIRC, LIHC and PRAD datasets containing both normal and tumor samples **(Supplementary Table 1)**. All genes were ranked by their differential expression based on their log_2_ fold changes. Query gene sets for GSEA included pathways from KEGG, Reactome, WikiPathways and MPath. GSEA results were filtered to include pathway gene sets with *p*-values below 0.05 and a minimum gene set size of 10 or a maximum gene size of 3000. Similarly, GSEApy was used to perform single sample GSEA (ssGSEA; Barbie *et al.*, 2009) **(Supplementary Table 1)** to acquire sample wise pathway scores using FPKM-UQ for BRCA, KIRC, LIHC, OV and PRAD datasets, irrespective of phenotype labels (Barbie *et al.*, 2009). Datasets were filtered to only include normalized expression data for genes found in the pathway gene sets of KEGG, Reactome, WikiPathways and MPath and then used for ssGSEA. Expression data were ranked and sample-wise Normalized Enrichment Scores were obtained.

#### 5.3.3. Pathway topology-based enrichment

To evaluate PT-based methods, we selected the well-known and highly-cited Signaling Pathway Impact Analysis (SPIA) method (Tarca *et al*., 2008) for two main reasons: firstly, the guidelines outlined by a comparative study on topology-based methods (Ihnatova *et al*., 2018) recommend the use of SPIA for datasets with properties similar to TCGA (i.e., possessing two well-defined classes, full expression profiles, many samples and numerous differentially expressed genes). Secondly, SPIA has been reported to have a high specificity whilst preserving dependency on topological information (Ihnatova *et al*., 2018). Because the R/Bioconductor’s SPIA package only contains KEGG pathways, we converted the pathway topologies from the three databases used in this work to a custom format in a similar fashion as graphite (Sales *et al.*, 2018) (**Supplementary Information)**. We declared significance for SPIA based pathway enrichment, if the Bonferroni corrected *p*-value was < 5%.

#### 5.3.4. Evaluation based on enrichment of pathway analogs

To better understand the impact of database choice, we compared the raw *p*-value rankings (i.e., before multiple testing correction) of pathways analogs across each possible pair of databases (i.e., in KEGG and Reactome, Reactome and WikiPathways, and WikiPathways and KEGG) and in each statistical enrichment analysis (i.e., hypergeometric test, GSEA, and SPIA) with the Wilcoxon signed rank test. It assessed the average rank difference of the pathway analogs and reported how significantly different the results were for each database pair. Importantly, we only tested statistical enrichment of the analogous pathways in order to avoid statistical biases due to differences in the size of pathway databases.

### 5.4. Machine Learning

ssGSEA was conducted to summarize the gene expression profile mapping to a particular pathway of interest within a given patient sample, hence resulting in a pathway activity profile for each patient. We then evaluated the different pathway resources with respect to three machine learning tasks:

1. Prediction of tumor vs. normal
2. Prediction of known tumor subtype
3. Prediction of overall survival

#### 5.4.1. Prediction of tumor vs. normal

The first task was to train and evaluate binary classifiers to predict normal versus tumor sample labels. This task was conducted for four of the five TCGA datasets (i.e., BRCA, KIRC, LIHC and PRAD) while OV, which only contains tumor samples, was omitted. We performed this classification using a commonly used elastic net penalized logistic regression model (Zou *et al.*, 2005). Prediction performance was evaluated via a 10 times repeated 10-fold stratified cross-validation. Importantly, tuning of elastic net hyper-parameters (*□_1_*, *□_2_* regularization parameters) was conducted within the cross-validation loop to avoid over-optimism (Molinaro *et al.*, 2005).

#### 5.4.2. Prediction of tumor subtype

The second task was to train and evaluate multi-label classifiers to predict tumor subtypes using sample-wise pathway activity scores generated from ssGSEA. This task was only conducted for the BRCA and PRAD datasets, similar to the work done by Lim *et al*. (2018), because the remaining three datasets included in this work lacked subtype information. From the five breast cancer subtypes present in the BRCA dataset by the PAM50 classification method (Sorlie *et al.*, 2001), we included four subtypes (i.e., 194 Basal samples, 82 Her2 samples, 567 LumA samples and 207 LumB samples). These four were selected as they constitute the agreed upon intrinsic breast cancer subtypes according to the 2015 St. Gallen Consensus Conference (Coates *et al.*, 2015) and are also recommended by the ESMO Clinical Practice Guidelines (Senkus *et al.*, 2015). For the PRAD dataset, evaluated subtypes included 151 ERG samples, 27 ETV1 samples, 14 ETV4 samples, 38 SPOP samples and 87 samples classified as other (Cancer Genome Atlas Research Network, 2014). Similar to the approach by Graudenzi *et al.*, support vector machines (SVMs) (Cortes and Vapnik, 1995) were used for subtype classification by implementing a one-versus-one strategy in which a single classifier is fit for each pair of class labels. This strategy transforms a multi-class classification problem into a set of binary classification problems. We again used a 10 times repeated 10-fold cross-validation scheme, and the soft margin parameter of the linear SVM was tuned within the cross-validation loop via a grid search. We assessed the multi-class classifier performance in terms of accuracy, precision and recall.

#### 5.4.3. Prediction of overall survival

The third task was to train and evaluate machine learning models to predict overall survival of cancer patients. For this purpose, a Cox proportional-hazards model with elastic net penalty was used (Tibshirani, 1997; Friedman *et al*., 2010). Prediction performance was evaluated on the basis of five TCGA datasets (i.e., BRCA, LIHC, KIRC, OV and PRAD) (Table 3) using the same 10 times repeated 10-fold nested cross-validation procedure as described before. The performance of the model was assessed by Harrell’s concordance index (c-index; Harrell *et al.*, 1982), which is an extension of the well known area under ROC curve for right censored time-to-event (here: death) data.

#### 5.4.4. Statistical assessment of database impact on prediction performance

To understand the degree to which the observed variability of AUC values, accuracies and c-indices could be explained by the actually used pathway resource, we conducted a two-way Analysis of Variance (ANOVA). The ANOVA model had the following form:

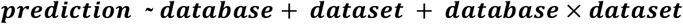

We then tested the significance of the database factor via an F-test. In addition, we performed Tukey’s post-hoc analysis to understand specific differences between databases in a dataset-dependent manner.

### 5.5. Software implementation

The workflow presented in this article consists of three major components: i) the acquisition and preprocessing of gene set and pathway databases, ii) the acquisition and preprocessing of experimental datasets and iii) the re-implementation or adaptation of existing analytical pipelines for benchmarking. We implemented these components in the `pathway_forte` Python package to facilitate the reproducibility of this work, the inclusion of additional gene set and pathway databases and to include additional experimental datasets.

The acquisition of KEGG, MSigDB, Reactome and WikiPathways was mediated by their corresponding Bio2BEL Python packages (Hoyt *et al.*, 2019; https://github.com/bio2bel) in order to provide uniform access to the underlying databases and to enable the reproduction of this work as they are updated. Each Bio2BEL package uses Python’s *entry points* to integrate in the previously mentioned ComPath framework in order to support uniform preprocessing and enable the integration of further pathway databases in the future, without changing any underlying code in the `pathway_forte` package. The network preprocessing defers to PathMe (Domingo-Fernández et al., 2019; https://github.com/pathwaymerger). Because it is based on PyBEL (Hoyt et al., 2018; https://github.com/pybel), it is extensible to the growing ecosystem of BEL-aware software.

While the acquisition and preprocessing of experimental datasets is currently limited to a subset of TCGA, it is extensible to further cancer-specific and other condition-specific datasets. We implemented independent preprocessing pipelines for several previously mentioned datasets (Table 3) using extensive manual curation, preparation and processing with the pandas Python package (McKinney, 2010; https://github.com/pandas-dev/pandas). Unlike the pathway databases, which were amenable to standardization, the preprocessing of each new dataset must be bespoke.

The re-implementation and adaptation of existing analytical methods for functional enrichment and prediction involved wrapping several existing analytical packages (Table 5) in order to make their application programming interfaces more user-friendly and to make the business logic of the benchmarking more elegantly reflected in the source code of `pathway_forte`. Each is independent and can be used with any combination of pathway database and dataset. Finally, all figures presented in this paper and complementary analyses can be generated and reproduced with the Jupyter notebooks located at https://github.com/pathwayforte/results/.

**Table 5.** Analytical packages wrapped by the `pathway_forte` package.

| Analysis | Package | Language | Source Code |
| --- | --- | --- | --- |
| Over-representation analysis | statsmodels | Python | https://github.com/statsmodels/statsmodels |
| GSEA | gseapy | Python | https://github.com/zqfang/gseapy |
| ssGSEA | gseapy | Python | https://github.com/zqfang/gseapy |
| Prediction tasks | scikit-learn | Python | https://github.com/scikit-learn/scikit-learn |
| Survival analysis | scikit-survival | Python | https://github.com/sebp/scikit-survival |
| SPIA | PyBEL-Tools<br>spia | Python<br>R | https://github.com/pybel/pybel-tools<br>https://bioconductor.org/packages/release/bioc/html/SPIA.html |

Ultimately, we wrapped each of these components in a command line interface (CLI) such that the results presented in each section of this work can be generated with a corresponding command following the guidelines described by Grüning *et al*. (2019). The scripts for generating the figures in this manuscript are not included in the main `pathway_forte`, but rather in their own repository within Jupyter notebooks at https://github.com/PathwayForte/results.

The source code of the `pathway_forte` Python package is available at https://github.com/PathwayForte/pathway-forte, its latest documentation can be found at https://pathwayforte.readthedocs.io and its distributions can be found on PyPI at https://pypi.org/project/pathway-forte.

The `pathway_forte` Python package has a tool chain consisting of pytest (https://github.com/pytest-dev/pytest) as a testing framework, coverage (https://github.com/nedbat/coveragepy) to assess testing coverage, sphinx (https://github.com/sphinx-doc/sphinx) to build documentation, flake8 (https://github.com/PyCQA/flake8) to enforce code and documentation quality, setuptools (https://github.com/pypa/setuptools) to build distributions, pyroma (https://github.com/regebro/pyroma) to enforce package metadata standards and tox (https://github.com/tox-dev/tox) as a build tool to facilitate the usage of each of these tools in a reproducible way. It leverages community and open source resources to improve its usability by using Travis-CI (https://travis-ci.com) as a continuous integration service, monitoring testing coverage with Codecov (https://codecov.io) and hosting its documentation on Read the Docs (https://readthedocs.org).

## Acknowledgements

The authors would like to thank Mohammad Asif Emon for his assistance in conducting SPIA and Jan-Eric Bökenkamp for his assistance in processing the TCGA datasets. Furthermore, we would like to thank Jonas Klees and Carina Steinborn for generating the visuals in this paper. Finally, we would like to thank the curators of KEGG, Reactome and WikiPathways as well as the TCGA network for generating the pathway content and datasets used in this work, respectively.

## Competing interests

H.F. received salaries from UCB Biosciences GmbH. UCB Biosciences GmbH had no influence on the content of this work.

## Funding

This work was supported by the EU/EFPIA Innovative Medicines Initiative Joint Undertaking under AETIONOMY [grant number 115568], resources of which are composed of financial contribution from the European Union’s Seventh Framework Programme (FP7/2007-2013) and EFPIA companies in kind contribution.

## Authors’ contributions

DDF conceived and designed the study. SM and DDF conducted the main analysis and implemented the Python package. HF supervised methodological aspects of the analysis. CTH and AG assisted technically in the analysis of the results. MHA acquired the funding. SM, HF, CTH, MHA and DDF wrote the paper.

## Notes

https://github.com/pathwayforte

