## Supplementary File for "The Impact of Pathway Database Choice on Statistical Enrichment Analysis and Predictive Modeling"

Supplement Material

| **Method** | **Parameters** |
| --- | --- |
| GSEA | method=ratio_of_classes  number of permutations=500  permutation_type=phenotype  min_size=15/max_size=3000 |
| ssGSEA | method=rank  min_size=15/max_size=3000 |

**Supplementary Table 1.** Parameters used in functional class enrichment methods.

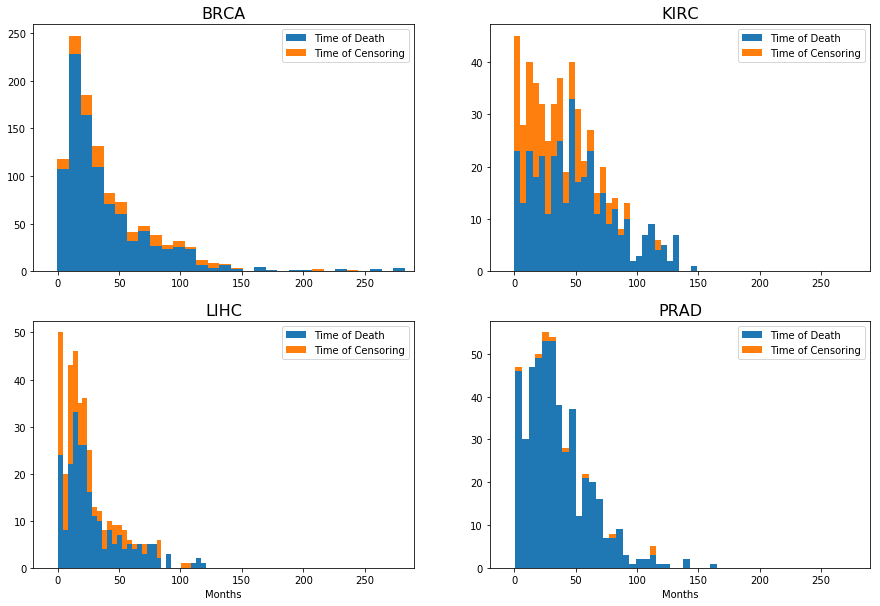

**Supplementary Figure 1.** Histogram of the lifetime data for four TCGA datasets used for survival analysis (i.e., BRCA, LIHC, KIRC and PRAD). This visualization can be reproduced using the following Jupyter notebook:<https://nbviewer.jupyter.org/github/pathwayforte/results/blob/master/notebooks/others/prediction/survival/survival_curves.ipynb>.

### **Adapting BEL for SPIA analysis**

This section outlines the transformation from the pathway BEL networks to SPIA. To conduct this transformation, we implemented a new module in the PathMe package called “spia”. Therefore the SPIA files can be regenerated in the future by running a simple command: “python -m pathme export spia” (assuming that the BEL files are already parsed). Similar to the graphite R package, PathMe processes the networks in a way that complex nodes (i.e., reactions and complexes) are flattened and proteins and RNAs are collapsed to genes as well as variants. For further details, we defer to the PathMe paper. Specifically for Reactome, we used its hierarchy to infer the networks of the pathways containing children. The resulting BEL networks are then converted into a connectivity matrix filled with 1s in the case that there exists a relationship between two nodes, or 0s otherwise. This connectivity matrix spans over multiple relationships (thus, has multiple dimensions) that are mapped directly from BEL to the SPIA custom format (see PyBEL-Tools SPIA module for more details).
